# Transcriptome assembly and annotation of johnsongrass (*Sorghum halepense*) rhizomes identifies candidate rhizome-specific genes

**DOI:** 10.1101/243956

**Authors:** Nathan Ryder, Kevin M. Dorn, Mark Huitsing, Micah Adams, Jeff Ploegstra, Lee DeHaan, Steve Larson, Nathan L. Tintle

## Abstract

Rhizomes facilitate the wintering and vegetative propagation of many perennial grasses. *Sorghum halepense* (johnsongrass) is an aggressive perennial grass that relies on a robust rhizome system to persist through winters and reproduce asexually from its rootstock nodes. This study aimed to sequence and assemble expressed transcripts within the johnsongrass rhizome. A *de novo* transcriptome assembly was generated from a single johnsongrass rhizome meristem tissue sample. A total of 141,176 probable protein-coding sequences from the assembly were identified and assigned gene ontology terms using Blast2GO. The johnsongrass assembly was compared to *Sorghum bicolor*, a related non-rhizomatous species, along with an assembly of similar rhizome tissue from the perennial grain crop *Thinopyrum intermedium*. The presence/absence analysis yielded a set of 259 johnsongrass contigs that are likely associated with rhizome development.

## Introduction

Rhizomes are the horizontally-aligned subterranean stems which allow a perennial plant to grow back from dormancy after a period of harsh seasonal conditions. With rhizomes at sufficient depth, an herbaceous plant may reemerge each year without having to make the extensive investment into root growth required by germination. The growing season for a rhizomatous plant is lengthened when its roots reach below the top layer of the soil, reducing the effects of environmental stresses such as temperature. Furthermore, a large persisting root network allows perennial plants to limit soil erosion, reduce runoff, and store more carbon underground as compared with an annual crop (Cox, Glover, Van Tassel, Cox, & DeHaan, 2006).

Annual species such as wheat or corn require frequent disruptions from tillage and can only achieve diminished root depth and length before harvest. Soils planted with annual crops are then susceptible to excessive erosion. A 100 year agricultural experiment revealed that plots with continuously cropped corn retained less than half the amount of topsoil as plots with only perennial grasses (Gantzer, Anderson, Thompson, & Brown, 1990). Annual root systems also exacerbate nutrient runoff. Corn, wheat, and rice have shown nitrogen fertilizer uptake efficiencies ranging from 18 to only 49% (Cassman, Dobermann, & Walters, 2002). In fact, the loss of nitrate N through subsurface drainage may be 30 to 50 times greater in annual than in perennial crops (Randall & Mulla, 2001).

Perennial plants present further ecological advantages via carbon sequestration. Typical annual crops increase soil organic carbon by 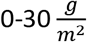 in a year, while perennials plants were found to accumulate 32-44 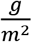 (Robertson, Paul, & Harwood, 2000). Perennial crops not only store more carbon, but also have no requirement to expend fossil fuels on tillage. They have a negative global warming potential, in CO_2_ equivalents, while annual crops actually increase atmospheric carbon (Robertson et al., 2000). *Sorghum halepense* is a perennial grass which has been classified as an invasive weed in 53 countries (Holm, Plucknett, Pancho, & Herberger, 1977). S. halepense rhizomes, which are known to regenerate even when cut into pieces, allow the plant to endure winters and extend locally (Howard, 2004). One plant may grow 275 feet of rhizomes in a single growing season. Moreover, johnsongrass is self-fertile with a high rate of seed production, yielding more than 80,000 seeds in the same growing season (Howard, 2004; Johnson, Kendig, Smeda, & Fishel, 1997). This propensity for reproduction, coupled with rapid growth relative to native grasses, can give johnsongrass a competitive edge and lead to monocultures in unsupervised areas. Johnsongrass also exhibits the ability to grow in a wide range of environmental conditions and has resistances to many common herbicides and pathogens (Howard, 2004; Johnson et al., 1997). With such vigorous root systems and high fecundity, this species is an excellent model for understanding one important strategy of the perennial growth habit. Johnsongrass rhizomes are likely to be useful for investigations of gene expression related to resource management and asexual reproduction. Thus, observations of highly expressed or rhizome-specific transcripts in S. halepense could improve prediction of perennial behavior in perennial grasses (Jang et al., 2009).

Attempts at breeding a perennial hybrid from diploid *S. bicolor* and tetraploid *S. halepense* began in earnest before 2003 at the Land Institute in Salina, Kansas (Cox, Glover, Van Tassel, Cox, & DeHaan, 2006). Developing a hybrid with sufficient yield may require an extensive time-frame, but this can be expedited through the use of genetically informed techniques. Marker-assisted selection could make use of QTL's discovered in johnsongrass that are linked to rhizome activity (Xiong et al., 2007).

Previous studies have connected johnsongrass RNA with rhizome function (Jang et al., 2006, 2009), but lack the sheer quantity of data expected from sequencing technologies today. This study provides a large, well-annotated transcriptome for rhizome tissue from johnsongrass. The sequenced rhizome RNA was assembled, filtered, and annotated with up-to-date tools for non-model organisms. A novel transcriptome for johnsongrass rhizomes then facilitates the discovery of additional rhizome-enriched transcripts. Further analysis with BLAST compared assembled johnsongrass sequences with non-rhizomatous and rhizomatous species, and has led to a reduced set of candidate genes for rhizome-related function.

## Materials and Methods

### Plant Origin and RNA Extraction

Johnsongrass plants were harvested at the coordinates 38°46’15.46” N and 97°34’21.77” W near Salina, Kansas on July 7, 2015. They were shipped overnight on dry ice to Sioux Center, Iowa where the rhizome apical meristems were removed, immersed in liquid nitrogen, and placed in a −80°C freezer. RNA was extracted from four samples of rhizome node buds using the RNeasy Plant Mini Kit (Qiagen) precisely following recommended protocols. DNase digestion was then performed with the TURBO DNA-*free* kit (Thermo Fisher Scientific), followed by the RNA cleanup procedure from the RNeasy mini handbook. Each sample received an A_260_/A_280_ ratio greater than 2 in a NanoDrop 2000 Spectrophotometer (Thermo Scientific). All four samples were then sent to the University of Minnesota Genomics Center for sequencing.

*Thinopyrum intermedium* rhizome tissue was obtained from a clone of a plant derived from The Land Institute’s breeding program (C3-3471). RNA was extracted as described above.

### Library Construction and Next Generation Sequencing

The four samples were run through a denaturing agarose gel to visualize RNA integrity. Two samples depicting intact RNA were pooled and submitted for library prep with the TruSeq Stranded mRNA Library Prep kit (Illumina). The resulting cDNA library was sequenced in a single lane of High Output (2x125 bp) on the Illumina HiSeq 2500 platform. The raw reads were trimmed in BBduk (Bushnell, 2014), removing adaptors and reads that were duplicated, short, or low quality.

*Thinopyrum intermedium* rhizome RNA libraries were prepared using the Illumina TruSeq RNA Library Prep Kit v2 by the University of Minnesota Genomics Center and sequenced across three lanes of an Illumina HiSeq 2000 (2x100 bp) run. Libraries were size-selected with a Caliper LabChip XT (PerkinElmer) to have an approximate insert size of 200 bp. A total of 5.7 Gigabases of high quality (>Q30) data was generated for this library (65,413,146 paired-end reads).

### Trinity Assembly and TransDecoder Identification of Protein Coding Sequences

The cleaned reads were *de novo* assembled in Trinity (version 2.4.0) (Grabherr et al., 2011; Haas et al., 2013). Next, the assembly was run through TransDecoder (version 3.0.1), a companion to Trinity which uses open reading frames within assembly transcripts to find likely coding regions (Haas & Papanicolaou, 2017).

### Annotation with Blast2GO PRO

The predicted coding sequences from TransDecoder were annotated in Blast2GO PRO, an all-in-one tool that befits large-scale functional annotation with Gene Ontology (GO), especially for an original assembly of a non-model organism. The suite performs the BLAST algorithm to align FASTA-formatted sequence inputs with homologs in specified databases. Annotations are mapped to query sequences by their association with BLAST hits (Conesa et al., 2005).

The BLASTX-fast search was used, aligning the coding sequences against the NCBI non-redundant protein database (NR). Query sequences with BLAST hits were mapped to GO terms and assigned functional annotations where available.

### Comparative Transcriptomics with Sorghum bicolor and Thinopyrum intermedium Rhizomes

Two sequenced genomes closely related to *Sorghum halepense* were selected: *Sorghum bicolor* (a close annual relative and possible ancestor of *S. halepense* (Jang et al., 2009)) and *Thinopyrum intermedium* (a recently sequenced rhizomatous perennial grass). The JGI Plant Flagship genome of non-rhizomatous *S. bicolor* (version 3.1.1) was accessed from the NCBI RefSeq database (Accession ID: ABXC03000000). The protein model reference sequences were formatted into a protein BLAST database. Predicted transcripts for the rhizomatous species *Thinopyrum intermedium* were obtained from a pre-publication annotated assembly (JGI annotation version 2.1) (Dorn et al, unpublished – pending release onto Phytozome) and were formatted into a second protein BLAST database for comparison.

Using the *S. halepense* proposed protein coding sequences as queries, two BLASTX searches were performed against the custom protein databases of *Th. intermedium* and *S. bicolor*. The query sequences that received one or more BLAST hit (e-value < 10^−20^) were cross-listed between the two searches, and removed. The *S. halepense* sequences that aligned in rhizomatous *Th. intermedium* but had no BLAST hits in non-rhizomatous *S. bicolor* were selected as candidates for rhizome-related function.

A rhizome-specific assembly for *Thinopyrum intermedium* was also generated in this study (NCBI Accession Number pending) and used to further reduce candidates. The assembly was aligned to the *Th. intermedium* protein BLAST database with a BLASTX search, identifying the sequences from rhizomes in the full *Th. intermedium* gene model set. The *Th. intermedium* predicted peptides declared present in rhizomes were then intersected with the *S. halepense* queries that had a *Th. intermedium* BLAST hit but not a hit from *S. bicolor*. The sequences that resulted are likely related to rhizome function, as they were expressed in *Th. intermedium* and *S. halepense* rhizomes, but do not appear in the genome of a closely related annual grain, *S. bicolor*.

## Results

### Assembly

Johnsongrass rhizome RNA was extracted and sequenced to yield 286,173,708 paired reads with an average Phred score > 30 (NCBI Sequence Read Archive SRX3427471). The reads were trimmed and assembled *de novo* in Trinity to produce 196,266 contigs. Using TransDecoder (version 3.0.1), 141,176 of the assembled contigs were identified as possible protein coding sequences. The proposed coding sequences have an average length of 825 base pairs (bp) and range from 297 to 15,174 bp. Figure 1 illustrates the distribution of contig lengths for the filtered assembly. An N50 value of 1,071 bp indicates that the contigs of this length or longer make up 50% of the total assembly length in base pairs. Table 1 lists the statistics for the sequencing/assembly process as a whole.

**Figure 1:**
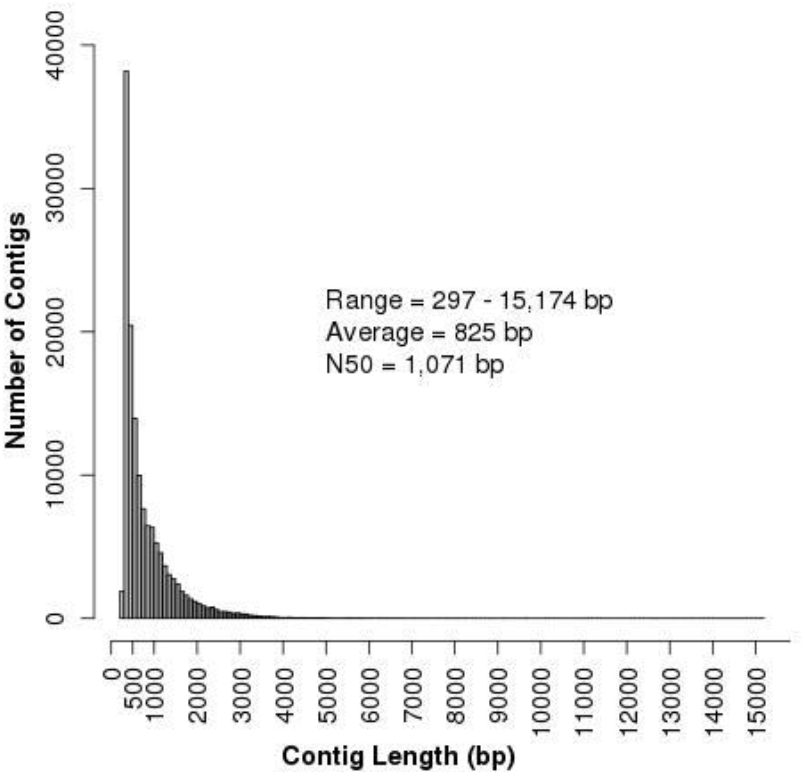
Distribution of base-pair lengths for contiguous sequences generated via Trinity assembly pipeline and predicted to be coding sequences in TransDecoder.

**Table 1:** Quantitative overview of sequenced reads before and after cleaning, assembled contigs, and predicted coding sequences from TransDecoder.

Sequencing and assembly statistics
|  |  |
| --- | --- |
| Total number of raw reads | 286,173,708 x 2 |
| Total number of cleaned reads | 283,167,847 x 2 |
| Total number of assembled contigs | 196,226 |
| Total number of assembled genes | 93,708 |
| Total number of protein coding sequences from TransDecoder | 141,176 |
| Mean length of coding sequences (bp) | 825 |
| Median length of coding sequences (bp) | 570 |
| N50 value of coding sequences (bp) | 1,071 |
| Range of coding sequence length (bp) | 297 - 15,174 |
| GC content | 53.37% |

### Annotation

A BLASTX-fast search against the NCBI non-redundant protein database (NR) aligned 126,750 (90%) of the johnsongrass predicted protein coding sequences with at least one hit in NR. GO terms associated with BLAST hits were then mapped to 106,537 (75%) of the query sequences. Lastly, functional annotations were found for 97,319 (69%) of the sequences (Table 2).

**Table 2:**
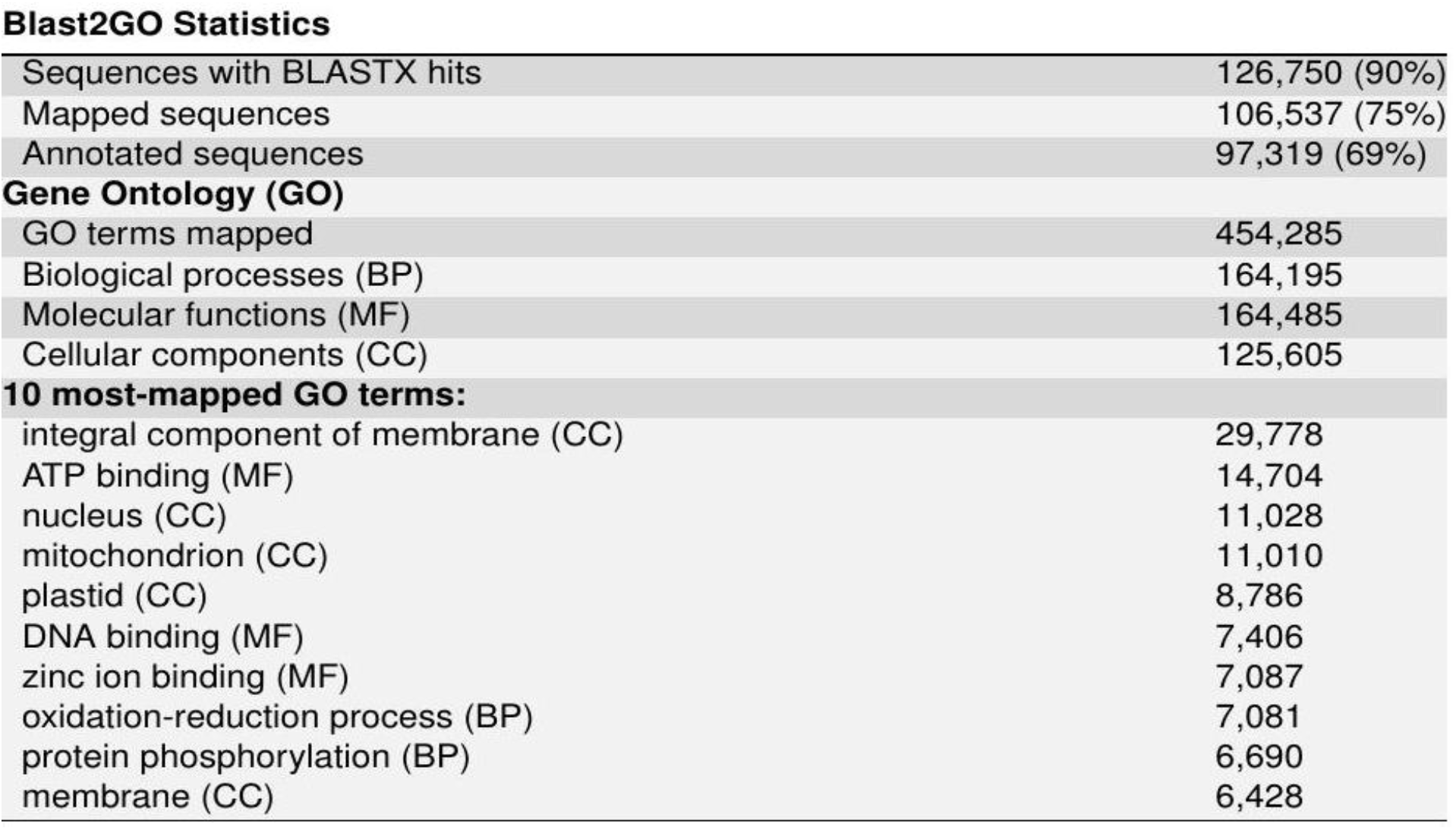
Results from the Blast2GO annotation suite.

*Sorghum bicolor* was the most prominently aligned species by far, receiving 96,303 of the top BLASTX hits (76%). The next highest amount of hits for a plant species was 7,801 (6%), from *Zea mays* (maize). A collection of fungi and bacteria accounted for over 13,000 (10%) of the top alignments, but many of these may be from symbiotes or pathogens dwelling within the roots.

**Figure 2:**
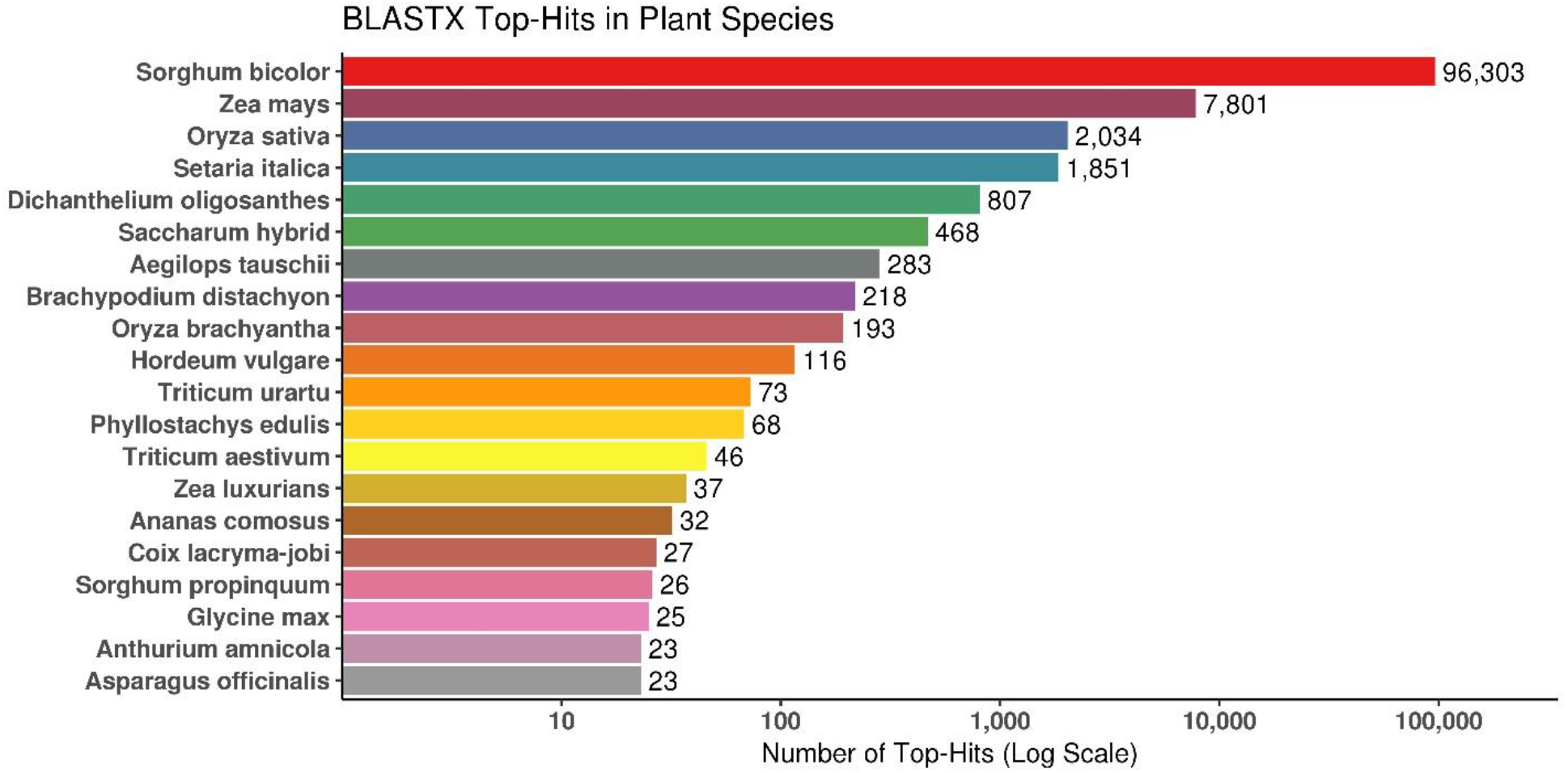
Species distribution of BLASTX top hits from the TransDecoder predicted coding sequences queried against the NCBI non-redundant protein database (NR). Only members of kingdom Plantae are included.

A total of 454,285 gene ontology terms were mapped to the johnsongrass contigs. These GO terms were split into three categories by their associations: 164,195 were for biological processes, 164,485 were for molecular functions, and 125,605 were for cellular components. The most-mapped term was associated with 29,778 sequences and was an “integral component of membrane” in cells. The results of gene ontology mapping and most common terms are listed in Table 2. Figure 3 depicts the distribution of GO terms at the second node of each directed acyclic graph generated in an annotation.

**Figure 3:**
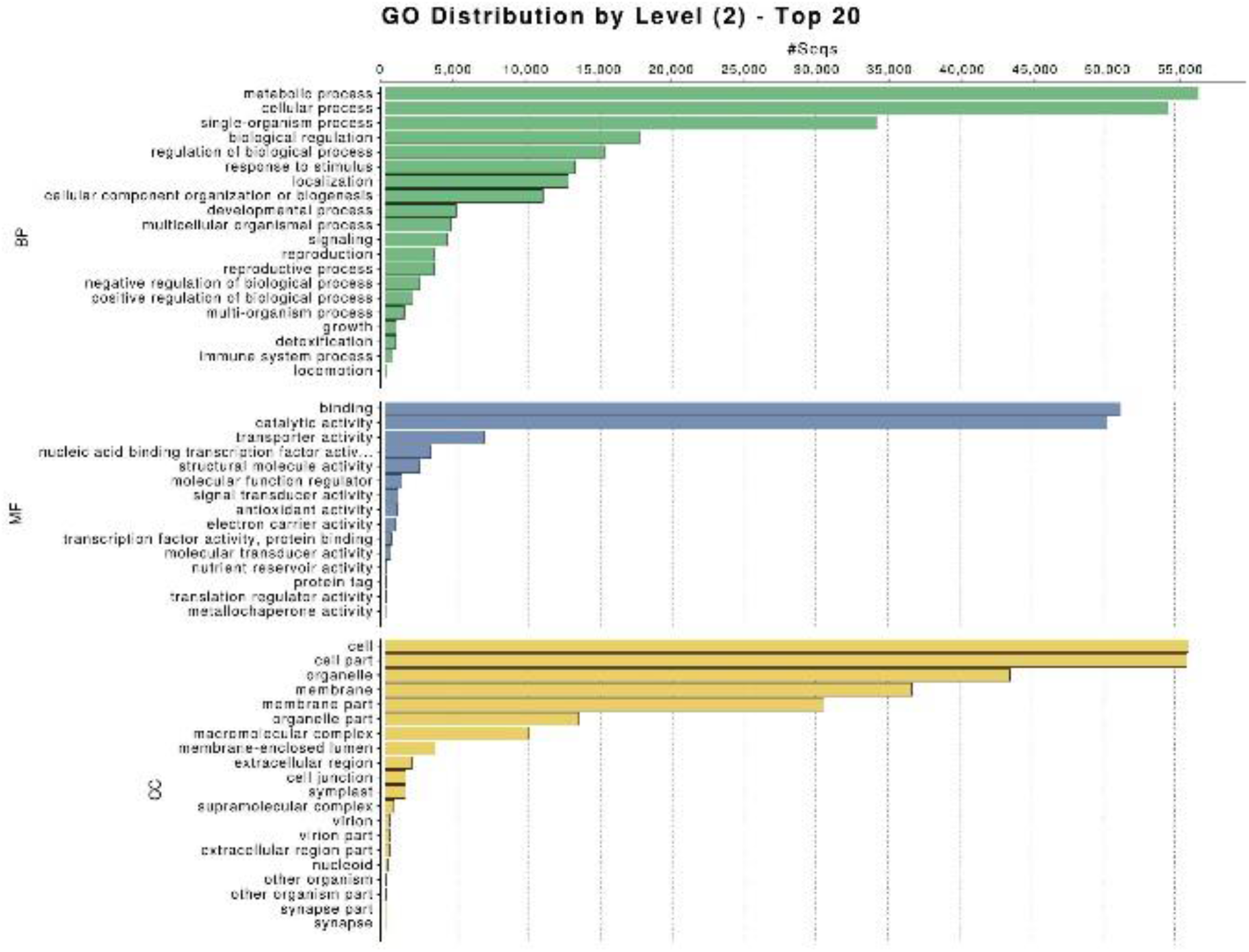
Distribution of GO level 2 terms by categories Biological Process (BP), Molecular Function (MF), and Cellular Component (CC). The top 20 mapped terms are listed for each category.

### Comparative Transcriptomics

Of the 141,176 johnsongrass coding sequences, only 991 of the johnsongrass contigs had a BLASTX hit from rhizomatous *Th. intermedium* but no hit from annual *S. bicolor*. Of these selected contigs, 259 had the same BLASTX hit from *Th. intermedium* as a sequence from the *Th. intermedium* rhizome assembly. Figure 4 depicts the BLASTX comparisons of the johnsongrass contigs. Table 3 lists the specific results of these searches as well as annotation results for the 259 candidate sequences.

**Table 3:** Overview of BLASTX comparisons of johnsongrass contigs with S. bicolor, Th. intermedium, and a rhizome assembly from Th. intermedium. Also including the most-mapped GO terms among the 259 candidate sequences.

BLASTX comparisons results
|  |  |
| --- | --- |
| Total number of *S. halepense* query sequences | 141,176 |
| Sequences with $\geq 1$ hit from *S. bicolor* | 105,365 |
| Sequences with $\geq 1$ hit from *T. intermedium* | 91,029 |
| Sequences with $\geq 1$ hit from both organisms | 90,038 |
| Sequences with no hits | 34,820 |
| Sequences with $\geq 1$ hit from **only** *S. bicolor* | 15,327 |
| Sequences with $\geq 1$ hit from **only** *T. intermedium* | 732 |
| Sequences with $\geq 1$ hit from **only** *T. intermedium* and in the rhizome assembly | 259 |
| **10 most-mapped GO terms in candidate sequences:** |  |
| integral component of membrane (CC) | 52 |
| mitochondrion (CC) | 44 |
| nucleic acid binding (MF) | 36 |
| RNA-directed DNA polymerase activity (MF) | 36 |
| RNA-dependent DNA biosynthetic process (BP) | 36 |
| membrane (CC) | 33 |
| plastid (CC) | 33 |
| nucleus (CC) | 22 |
| proteolysis (BP) | 18 |
| RNA binding (MF) | 17 |

**Figure 4:**
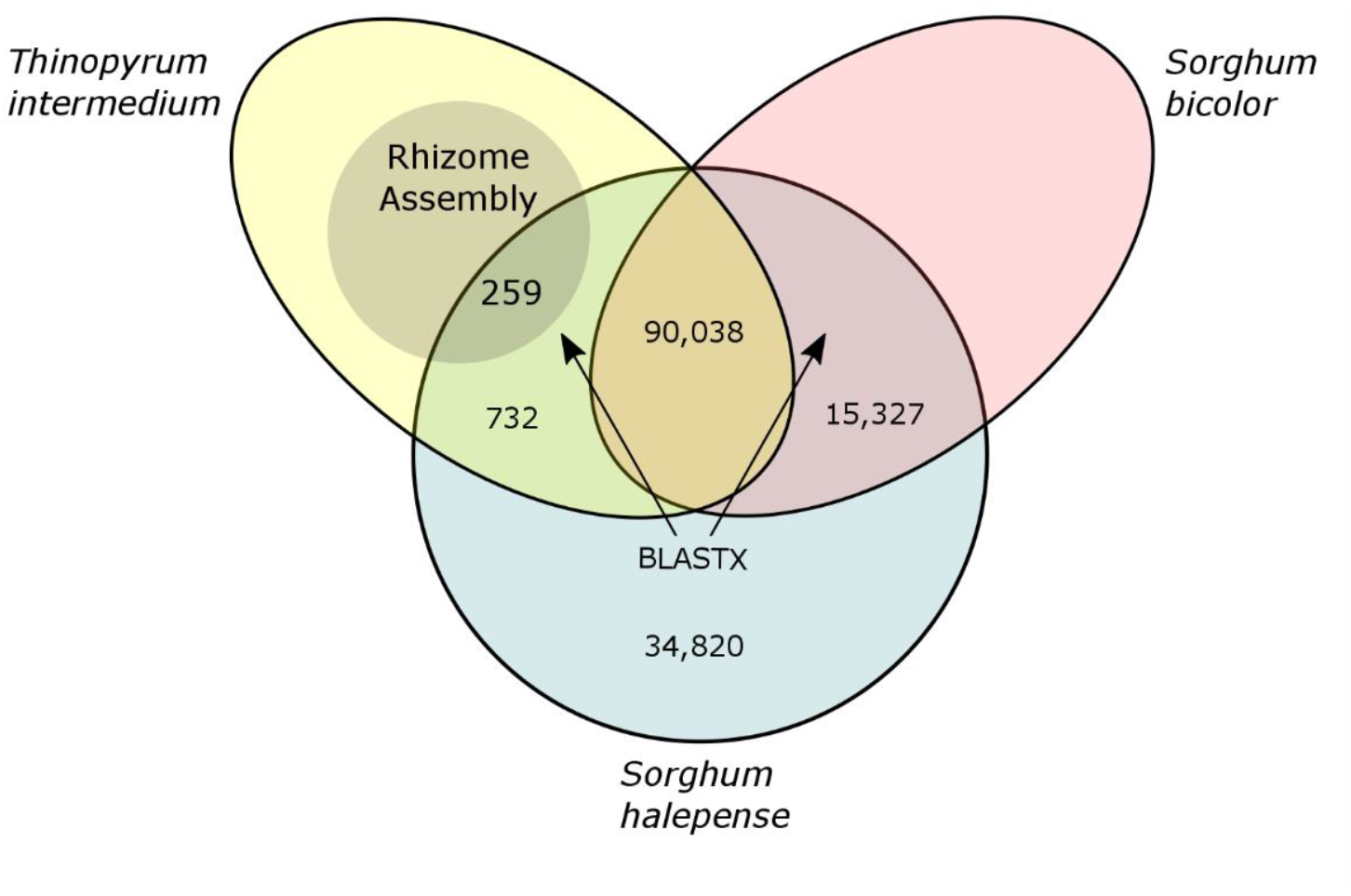
Sorghum halepense transcriptome comparison with S. bicolor and Th. intermedium. A Venn diagram depicts the distribution of S. halepense coding sequences as they have BLASTX hits against Th. intermedium, S. bicolor, both, or neither. An assembly of rhizome RNA from Th. intermedium was cross-listed with the S. halepense sequences that had BLASTX hits from Th. intermedium but not S. bicolor. All quantities refer to unique S. halepense contigs.

## Discussion/Conclusion

The identification of genes essential to perenniality remains an important, and yet, elusive biological problem, especially in efforts to perennialize grain crops and enhance the sustainability of food production. Recent efforts to gather expression data on rhizomatous tissue are providing important lists of candidate genes to accelerate the process of uncovering the biological pathways related to perenniality. This study provides an important step in the process, as well as providing a larger set of contigs (141,176 predicted contigs) for the, as yet, unsequenced *Sorghum halepense*.

Using presence/absence analysis via BLASTX searches against a *Th. intermedium* rhizome assembly and other genomes, 259 *S. halepense* sequences were selected as potentially related to rhizome behavior. These 259 candidate sequences will now be the focus of further bioinformatics analysis (e.g. protein functional prediction, enhanced annotation, and BLASTing). Especially promising candidates may become the target of genetic modification (e.g., knockout; over-expression) experiments to further elucidate genes critically part of grain crop perenniality.

Quantitative trait loci associated with rhizome function may be used to improve breeding practices for perennial traits in a Sorghum hybrid. This study has produced a comprehensive dataset which facilitates further research with *S. halepense* and other perennial grains.

## Data Accession

The johnsongrass dataset has been registered with NCBI as a BioProject (PRJNA417857). The raw read files are available in the Sequence Read Archive (SRP125786), and this Transcriptome Shotgun Assembly project has been deposited at DDBJ/EMBL/GenBank under the accession GGDZ00000000. The version described in this paper is the first version, GGDZ01000000. The results from Blast2GO, in .b2g or .txt file formats, are stored at <pending>.

The *Thinopyrum intermedium* dataset has been registered with NCBI as a BioProject (PRJNA428355).

The raw read files are available in the Sequence Read Archive (SRP127982), and this Transcriptome Shotgun Assembly project has been deposited at DDBJ/EMBL/GenBank under the accession <pending>. The version described in this paper is the first version, <pending>.

